# Water lily (*Nymphaea thermarum*) draft genome reveals variable genomic signatures of ancient vascular cambium losses

**DOI:** 10.1101/2019.12.18.881573

**Authors:** Rebecca A. Povilus, Jeffery M. DaCosta, Christopher Grassa, Prasad R. V. Satyaki, Morgan Moeglein, Johan Jaenisch, Zhenxiang Xi, Sarah Mathews, Mary Gehring, Charles C. Davis, William E. Friedman

## Abstract

For more than 225 million years, all seed plants were woody trees, shrubs, or vines (1–4). Shortly after the origin of angiosperms ~135 million years ago (MYA) (5), the Nymphaeales (water lilies) became one of the first lineages to deviate from their ancestral, woody habit by losing the vascular cambium (6), the meristematic population of cells that produces secondary xylem (wood) and phloem. Many of the genes and gene families that regulate differentiation of secondary tissues also regulate the differentiation of primary xylem and phloem (7–9), which are produced by apical meristems and retained in nearly all seed plants. Here we sequence and assemble a draft genome of the water lily *Nymphaea thermarum*, an emerging system for the study of early flowering plant evolution, and compare it to genomes from other cambium-bearing and cambium-less lineages (like monocots and *Nelumbo*). This reveals lineage-specific patterns of gene loss and divergence. *Nymphaea* is characterized by a significant contraction of the HD-ZIP III transcription factors, specifically loss of *REVOLUTA*, which influences cambial activity in other angiosperms. We also find the *Nymphaea* and monocot copies of cambium-associated CLE signaling peptides display unique substitutions at otherwise highly conserved amino acids. *Nelumbo* displays no obvious divergence in cambium-associated genes. The divergent genomic signatures of convergent vascular cambium loss reveals that even pleiotropic genes can exhibit unique divergence patterns in association with independent trait loss events. Our results shed light on the evolution of herbaceousness – one of the key biological innovations associated with the earliest phases of angiosperm evolution.

**Significance Statement:** For ~225 million years, all seed plants were woody trees, shrubs, or vines. Shortly after the origin of flowering plants ~135 million years ago, Nymphaeales (water lilies) became one of the first seed plant lineages to become herbaceous through loss of the meristematic cell population known as the vascular cambium. We sequence and assemble the draft genome of the water lily *Nymphaea thermarum*, and compare it to genomes of other plants that have retained or lost the vascular cambium. By using both genome-wide and candidate-gene analysis, we find lineage-specific patterns of gene loss and divergence associated with cambium loss. Our reveal divergent genomic signatures of convergent trait loss in a system characterized by complex gene-trait relationships.

## Introduction

In all seed plants, vascular tissue (xylem and phloem) initially develops in distinct bundles near apical meristems as part of primary growth and differentiation. In stems of most seed plants, the subsequent activation of a population of meristematic cells within and between the bundles forms a continuous layer of vascular cambium, which then produces rings of secondary xylem and phloem responsible for increases in stem or root girth [Figure 1] (9). Secondary growth provides the additional fluid transport capacity and mechanical support necessary for large, photosynthetic canopies to operate. A vascular cambium and production of secondary xylem and phloem are plesiomorphic for angiosperms. Loss of a vascular cambium is rare(1–3, 6) – even most herbaceous taxa (including *Arabidopsis)* form a vascular cambium at some point in their ontogenies. Only five losses of vascular cambium have been documented across angiosperms (7) (Nymphaeales, *Ceratophyllum*, monocots, *Nelumbo*, and Podostemaceae), which spans ~140 million years (5), ~350,000 species (10), and a diverse array of morphological adaptations and growth habits [Figure 1A].

**Figure 1:**
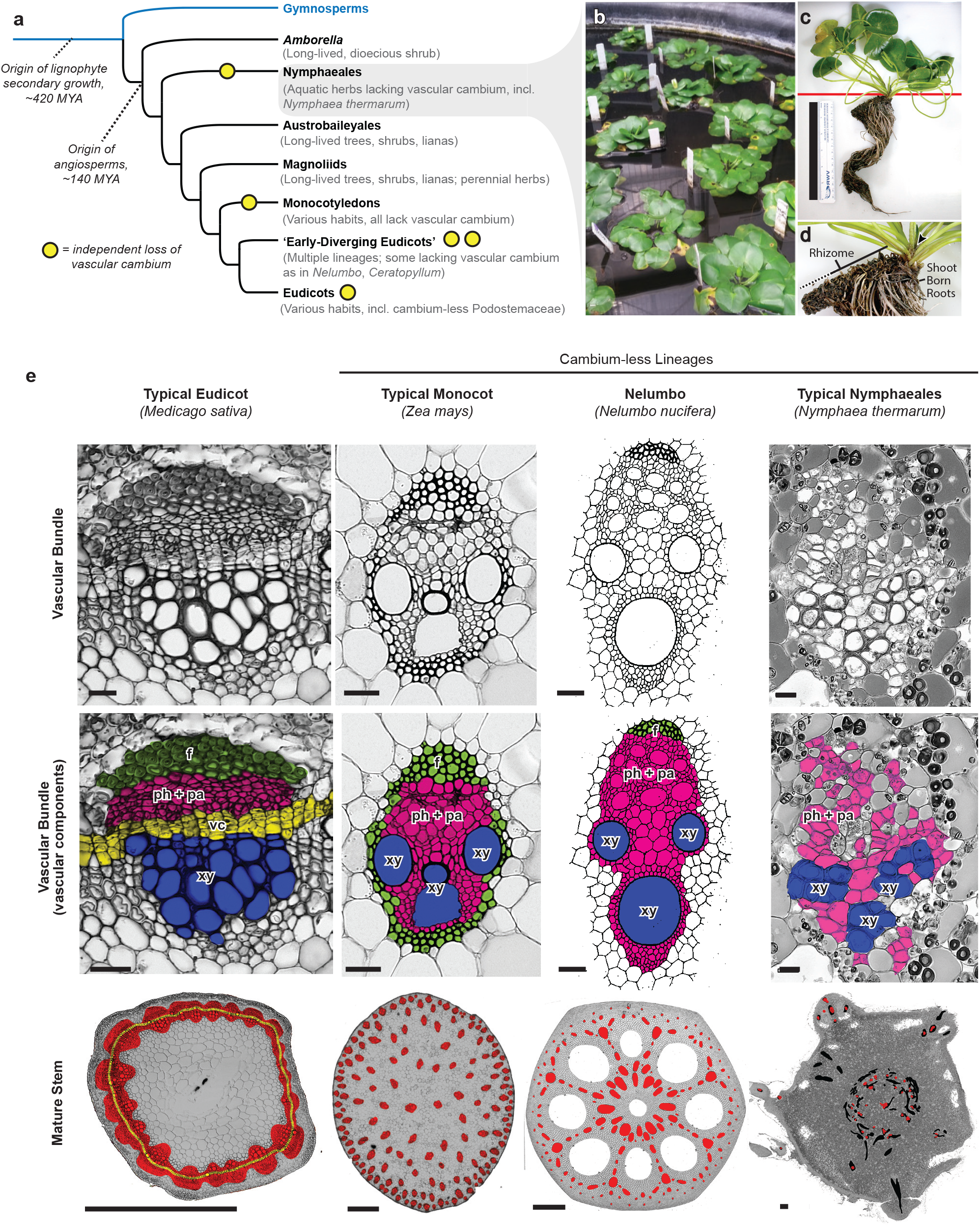
Vascular cambium evolution and stem vascular anatomy in flowering plants. A) Abbreviated phylogenetic relationships of major clades of seed plants, with notes on diversity of growth habits found in each group. B) Growth habit of *Nymphaea thermarum.* C) One individual of *N. thermarum.* Red line indicates in situ soil level. Scale bar = 15 cm. D) Close-up of rhizome from C). Arrowhead = apical meristem, black line = living rhizome section, dashed line = dead rhizome section. E) Stem vascular anatomy in clades of flowering plants. Cross sections were taken from maturing *(Medicago)* or mature *(Zea, Nelumbo, Nymphaea)* stems. Top row: micrographs *(Medicago, Zea, Nymphaea)* or drawings based on prepared slides (Nelumbo ((30)) of vascular bundles. Scale bars = 20 μm. Middle row: vascular components labeled and color-coded. ph + pa (pink) = phloem and vascular parenchyma, vc (yellow) = vascular cambium, f (green) = fibers, xy (blue) = xylem. Bottom row: whole stem sections. red = vascular bundles and tissue in cross section, yellow = vascular cambium, black = vascular bundles that are part of a leaf or flower trace, grey = non-vascular tissue. Scale bars = 0.5 mm.

Although primary and secondary vascular tissues in seed plants are derived from different meristematic cell populations, both are produced by precise patterning of the same suite of cell types. Accordingly, some of the same gene families or genetic modules function in both developmental contexts (7, 9, 11, 12): the CLAVATA (CLE)–WUSCHEL (WOX) regulatory loop promotes meristem identity at the cost of xylem formation (13–16), while multiple HD-ZIP III family members promote xylem differentiation and bipolar patterning of daughter cell types (17–21) *[SI Appendix, Figure* S1]. The functions of HD-ZIP III members and CLE-WOX modules have been documented during primary and secondary vascular differentiation in distantly related eudicots, including *Arabidopsis thaliana, Populus trichocarpa*, and *Zinnia elegans*, using loss-of-function and/or over-expression studies (8, 9, 22, 23). Whereas at least partial expression or functional redundancy between gene family members is common (18, 21), subfamilies of HD-ZIP III and CLE genes display ontogenic specialization.

The complexity of vascular differentiation and its regulation therefore raises the question of how relevant regulatory genes might evolve (24, 25) in association with the loss of the vascular cambium and secondary tissuse. Convergent loss or divergence of relevant genes has been documented in cases of convergent gene loss (26–28), suggesting a consistent, predictable association between trait loss and evolution of relevant genetic components. We use the repeated loss of the vascular cambum in angiosperms to test this association in a system characterized by pleiotropy and genetic redundancy.

## Results

The loss of vascular cambium during the early evolution of the water lily lineage (Nymphaeales) likely represents one of the oldest vascular cambium losses among seed plants. Stem vascular structure is sparsely documented within the Nymphaeales (29) and undescribed in *N. thermarum*. We investigated rhizome (underground stem) anatomy in *N. thermarum* and found no evidence of a vascular cambium or secondary vascular tissues [Figure 1D]. Additionally, *N. thermarum* vascular bundles contained no fibers, which differs from other cambium-less lineages (30) [Figure 1E]. Thus, all vascular tissues in *N. thermarum* are primary tissues derived from apical meristems.

Genomes for representatives of two of the five seed plant lineages that have independently lost the vascular cambium (monocots and *Nelumbo*) are already available. To create a genomic resource for *N. thermarum*, we combined data generated from short insert libraries (~40× coverage) and mate-pair libraries (~20× coverage) to create a draft genome assembly *[SI Appendix*,Table S1]. The total length of the assembled genome size was 368,014,730 bp after removing scaffolds < 1 kb, and a £-mer analysis of short insert reads estimated the genome size at 497,339,103 bp. These values are ~74% and ~100%, respectively, of the 1C size estimate of 0.51 pg as evaluated by flow cytometry (31) (roughly 498,780,000 bp). We also generated and assembled transcriptomes from leaves, roots, and reproductive material. When combined with available plant protein data, we annotated 25,760 protein-coding genes that overlapped input transcript or protein evidence, a Pfam domain, and/or a hit to the UniProt/SwissProt database [SI *Appendix*, Figure S2]. By comparison, the genome of *Amborella* (32) is annotated with 26,846 protein-coding genes [SI *Appendix*, Figure S2]. The *N. thermarum* assembly contained 865/956 (90%) of conserved plant proteins (33), suggesting that its genic regions were well represented. Whereas changes in non-coding regions are likely important for morphological evolution (28), we limited our analyses to coding regions due to higher confidence in their recovery and annotation across multiple genomes.

To detect significant contraction of gene families in cambium-less lineages, we first identified ortholog clusters from 28 genomes of seed plants, including three of five lineages that have independently lost the vascular cambium (i.e., *Nymphaea*, stem monocot, and *Nelumbo).* We estimated gene trees for 1,439 clusters that passed stringent filtering criteria (34), and then used them to estimate species trees with both concatenation and coalescent methods, including different subsets of the data (i.e.: fast-sites, slow-evolving sites, or all sites). All method and dataset combinations produced species trees with similar, strongly-supported topologies for relationships of interest, including support for *Amborella* as sister to all other extant angiosperm lineages [SI *Appendix*, Figure S3]. We used CAFÉ (35) to detect significant ortholog cluster contraction and expansion across all branches of our species phylogeny on a fossil-calibrated species tree (ASTRAL-II, all rate classes dataset) and the set of gene clusters sizes [Figure 2A, *SI Appendix*, Figure S4]. Among a total of 8,147 ortholog clusters included in the analysis, we found that 28 clusters underwent significant expansion within the Nymphaeales, while 30 significantly contracted [SI *Appendix,Table* S2, *SI Appendix*, Figure S4]. Few clusters were contracted in more than one cambium-less lineage [Figure 2B]. Clusters expanded in *N. thermarum* were enriched for GO terms related to secondary metabolism (including carbohydrate and polysaccharide metabolism), biosynthesis, and response to stimuli. Contracted clusters in *Nymphaea* were most strongly enriched for nucleosome assembly, fatty acid biosynthetic process, response to auxin stimulus, and meristem initiation [Figure 2B, *SI Appendix*, Figure S5, *SI Appendix,Table* S3].

**Figure 2:**
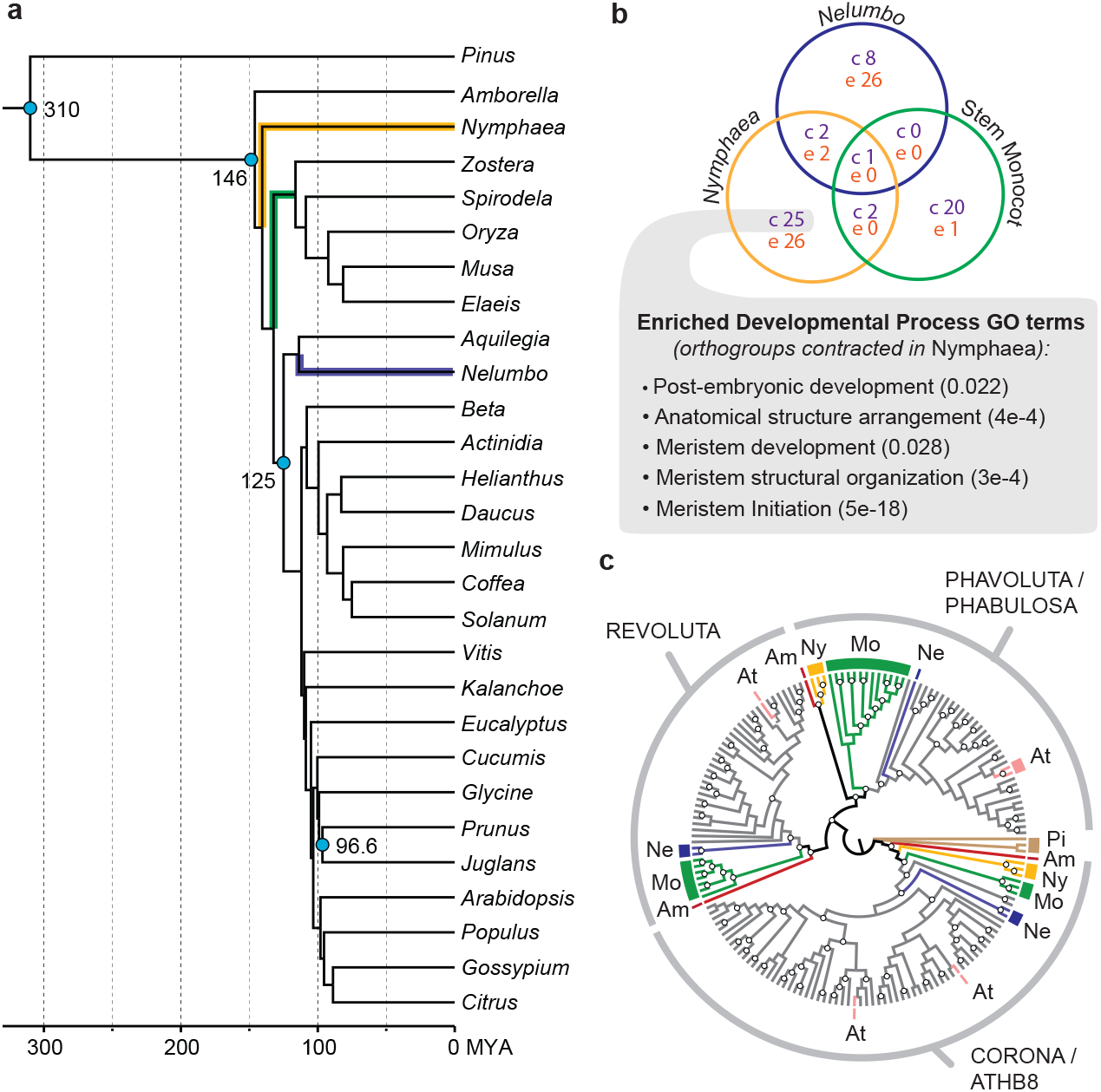
Gene family contraction in cambium-less lineages. A) Fossil-calibrated phylogenomic analysis, including representatives that have independently lost the vascular cambium: Nymphaea (yellow), monocots (stem lineage, green), and *Nelumbo* (indigo). Blue dots indicate age and placement of fossils or other estimates used for calibration. B) Venn diagram showing number of significantly contracted gene families in each cambium-less lineage. The set of gene families uniquely contracted in Nymphaea is enriched for “developmental process” GO terms related to meristem development and function (enrichment test p-value in parenthesis). C) Phylogeny of HD-zip III genes, with branches colored to reflect identity of genes from major lineages of seed plants (complete tree in Extended Figure 6). White dots denote branches with > 70% bootstrap support. Am = *Amborella trichopoda;* At = *Arabidopsis thaliana;* Mo = monocots; Ne = *Nelumbo nucifera;* Ny = Nymphaeaceae; Pi = *Pinus taeda.*

The contracted cluster associated with the meristem initiation GO term comprises Class III HD-ZIP genes, a gene family known to regulate differentiation of both primary and secondary vascular tissues (36, 37). Specifically, *N. thermarum* lacks a copy that belongs to the *REVOLUTA (REV)* subgroup [Figure 2C, *SI Appendix*, Figure S6]. We further determined that no *REV* homolog was present in the transcriptomes of several *N. thermarum* tissues, nor in an EST database of *Nuphar advena* (38), another member of the Nymphaeales [Figure 2C, *SI Appendix*, Figure S6]. The presence of a *REV* gene in *Amborella* and many other angiosperms [Figure 2C, *SI Appendix*, Figure S6] suggests that the lack of *REV* in *Nymphaea* and *Nuphar* represents a loss within the Nymphaeales. While some *REV* functions overlap with those of other HD-ZIP III members in embryogenesis and primary growth, *REV* mis-expression is known to impact both cambial initiation and xylary fiber differentiation during secondary growth in stems of *A. thaliana* and *P. trichocarpa* (17, 18, 21, 39). The combined absence of xylary fibers, vascular cambium initiation, and *REV* in *N. thermarum* [Figure 1D, Figure 2C, *SI Appendix*, Figure S6] is consistent with the hypothesis that *REV* is a key regulator of vascular cambium activity and differentiation of secondary tissues throughout angiosperms.

We further found that significant contraction of the HD-ZIP III family occurred along only two other branches [Figure 2A, *SI Appendix,Table* S2]: the terminal braches leading to *Zostera* (sea grass) and *Beta* (beet). These species are characterized by either the absence of secondary tissues *(Zostera)*, or by a highly anomalous pattern of vascular cambium activity that does not produce continuous rings of secondary xylem and phloem *(Beta)* (40–42). In contrast, some cambium-less lineages (some monocots and *Nelumbo)* retain homologs from all HD-ZIP III subfamilies [Figure 2C, *SI Appendix*, Figure S6]. Therefore, while all lineages with a significant contraction of the HD-ZIP III gene family display a loss of normal vascular cambium development, loss of HD-ZIP III genes are not necessary for vascular cambium loss in flowering plants. This result demonstrates variability in the patterns of gene loss associated with independent, homoplasious evolutionary events.

In addition to corroborating the importance of HD-ZIP III genes in vascular development, the ortholog clusters that are contracted in cambium-less lineages can be used to identify candidate genes for vascular cambium regulation. The set of clusters contracted in *N. thermarum* include the auxin-responsive SAUR proteins (43) and the BIM family of BES1-interacting proteins. Intriguingly, BIM and BES1 proteins regulate SAUR gene expression in other developmental contexts (44), and BES1 is known to regulate secondary growth (9) [SI *Appendix*, Figure S1]. Together with the contraction of the BIM and SAUR gene families in a species that lacks a vascular cambium, this suggests that BIM and SAUR genes may constitute a novel component of vascular cambium regulation. This remains to be experimentally tested.

The ortholog cluster expansion/contraction analysis, however, did not include all gene families of interest. The CLE (CLAVATA3/EMBRYO SURROUNDING REGION-RELATED) family of short (~21 amino acids) signaling peptides are known to regulate both primary and secondary tissue differentiation in flowering plants (45), but low sequence conservation among this gene family precludes many types of analysis. We therefore used CLANS clustering to assess patterns of relatedness between all putative CLE homologs (13) from Phytozome v12 (46) genomes, genomes of *N. thermarum* and *Nelumbo nucifera(47)*, available angiosperm transcriptomes from the OneKP initiative(48), and *N. thermarum* transcriptomes [Figure 3A] [SI *Appendix*, Figure S7]. We recovered major groups similar to results from previous studies, including a distinct CLE 41/42/44 group, called Group 2 (13, 49). Group 2 is associated with vascular differentiation and patterning (50), while the other groups are associated with root nodulation (Group 3) or a wide range of developmental processes, including the activity of shoot and root apical meristems (Group 1) (49). After adjusting for evolutionary distance of comparisons [SI *Appendix*, Figure S8] to calculate APAV (adjusted pairwise attraction values, scaled from 0 to 1 with 1 as the highest sequence similiarity), Group 2 amino acid sequences from species without a vascular cambium (NVC) and species with a vascular cambium (VC) were overall less similar than sequences from any two species that retain the vascular cambium (VC-VC) [Figure 3B]. This relationship was only present and significant (t-test p-value < 0.05 and a > 1% difference) for comparisons within Group 2 [SI *Appendix, Table* S4]. Therefore, divergence of CLE sequences from cambium-less taxa has occurred within Group 2, but not within the other CLE groups that are associated with primary root and shoot development.

**Figure 3:**
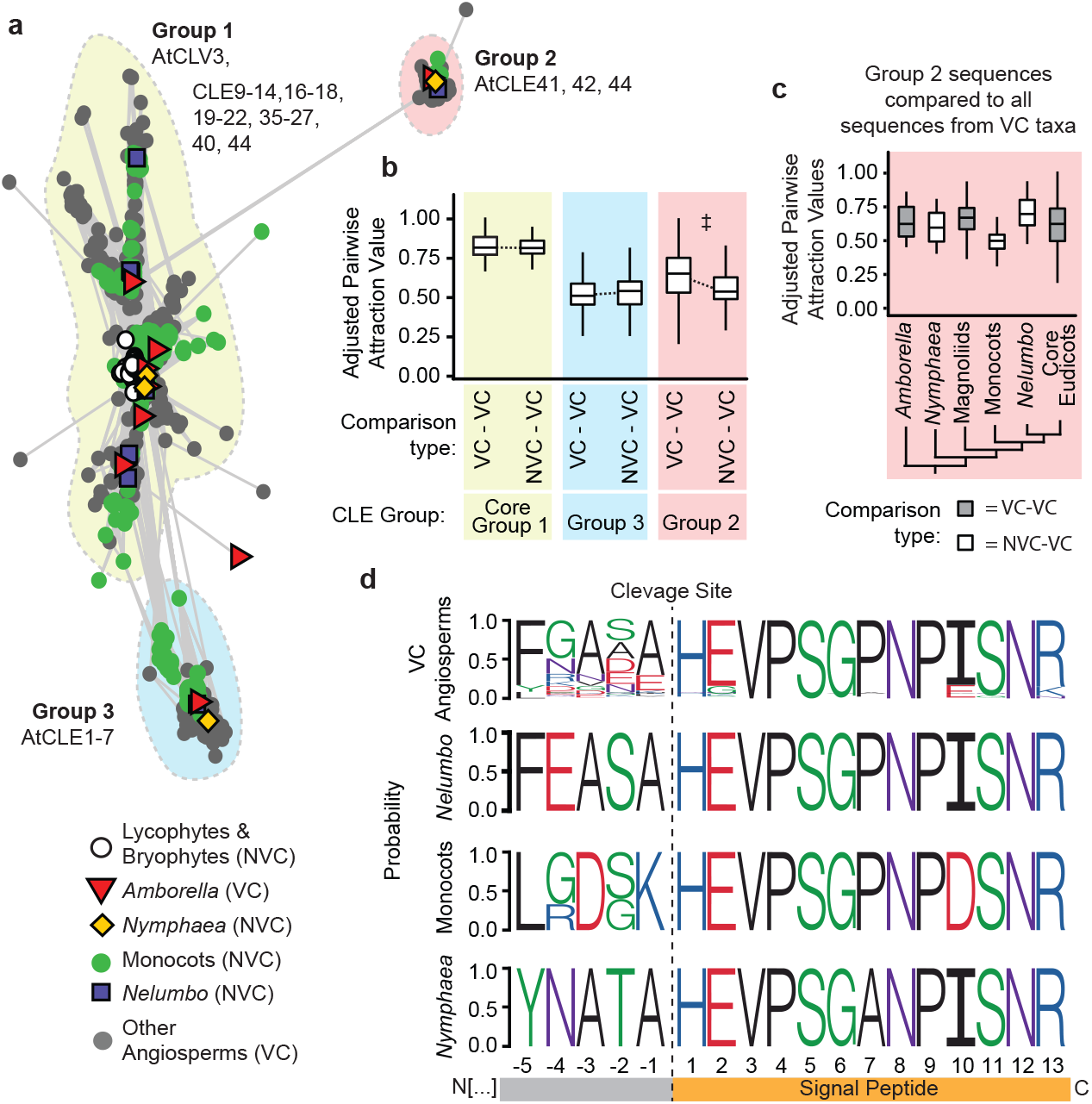
Evolution of vascular-development associated CLE peptides. A) 2-dimensional CLANS clustering of conserved amino acid sequences of CLE peptides in land plants. Dots/shapes represent individual sequences, coded to indicate whether they are from species within clades that have vascular cambia (VC) or do not have vascular cambia (NVC). Lines indicate PAV < 1e-5. B) APAV for all pairwise comparisons of CLE sequences between two VC taxa (VC-VC) or between one NVC and one VC taxa (NVC-VC), for each group of CLE peptides. Higher APAV indicate higher sequence similarity. ‡ indicates t-test adjusted p-value < 0.05 and median APAV difference > 1 % (Extended Data Table 1). C) APAV values for all sequences within Group 2 CLE peptides, for comparisons between all VC taxa and taxa within particular clades of flowering plants. D) Conserved peptide region of Group 2, including the 13 amino signal peptide at the C terminus and 5 amino acids upstream of the peptide cleavage site. For boxplots: center line, median; box limits, upper and lower quartiles; whiskers, 1.5x interquartile range; outliers removed from plots for clarity, but not from analysis.

The low similarity of Group 2 NVC-VC comparisons was largely driven by the divergence of monocot sequences [Figure 3C]. However, comparison of the C-terminal residues [Figure 3D] revealed that both monocot and *N. thermarum* sequences show lineage-specific divergences of the five amino acids that precede the signaling peptide. These residues are immediately adjacent to a cleavage site. Changes to residues surrounding the cleavage site may impact cleavage, and therefore mobility, of the peptide (49). Furthermore, in the *N. thermarum* sequence, a proline (P7) that is conserved across almost all angiosperm CLE peptides (49) is substituted with an alanine. Hydroxylation of this proline (as well as P4) is required for maturation of active Group 2 CLE peptides in *Arabidopsis* (51, 52). Additionally, prolines are necessary for certain types of kink formation (53); a substitution with alanine could negatively impact the ability of the peptide to conform to a receptor binding pocket (52, 54). In monocot sequences, a non-polar C-terminal binding anchor site (I9) is replaced with a negatively charged residue (D9). Intriguingly, co-option of Group 1 CLE peptides, but not Group 2, has been associated with secondarily-derived vascular cambia within monocots (55). This supports our conclusion that the observed divergence of Group 2 CLE peptides within some cambium-less taxa might be rendering them non-functional during vascular cambium development and activity.

## Discussion

Studies of convergent trait loss often document convergent patterns of sequence loss/divergence(26–28). However, many clear examples of gene loss in association with trait loss involve genes with one or few functions (56–58) and highly inter-connected genes are less likely to be lost or modified (59). We find, however, that *Nymphaea* is characterized by both divergence of a highly conserved CLE amino acid position and a significant contraction of the HD-ZIP III gene family of transcription factors — specifically loss of *REVOLUTA*, the HD-ZIP III member most closely associated with cambial activity and xylary fiber production in other angiosperms. In contrast, other lineages display unique signatures of cambium loss: monocot taxa all displayed a shared divergence of cambium-associated CLE signaling peptides, supporting a single loss of the cambium early in monocot evolution. Meanwhile, *Nelumbo* displays little obvious evidence of gene loss or divergence of key vascular cambium regulators. Our results confirm that not only can modification of pleiotropic genes occur in systems characterized by complex gene-trait relationships involving pleiotropy and genetic redundancy, but that genomic signatures of trait loss vary between lineages that represent homoplasious loss events.

Loss of the vascular cambium is highly correlated to transitioning from a terrestrial to an aquatic habit: Nymphaeales, *Nelumbo, Ceratophyllum*, and Podostemaceae are all aquatic lineages (60). Even in the case of monocots, the last common ancestor of the lineage has been proposed to have been semi-aquatic (61). Although there are many types of aquatic habits, submersion in or proximity to water means that aquatic plants typically do not require extensive mechanical reinforcement or high fluid transport capacity. Thus, the loss of selection for the functions performed by the vascular cambium appears to readily lead to the loss of vascular cambium formation. The variable genetic signatures of cambium loss that we document suggests that the evolution of genetic components of vascular cambium regulation is less predictable than the loss of the cambium itself.

The exploration of new growth habits, such as non-woody aquatic herbaceousness in Nymphaeales, is a hallmark of early angiosperm evolution (6). Yet, the genetic basis for the evolutionary diversification of early angiosperms has remained largely inaccessible due to the lack of a tractable system for genetic analyses outside of monocots or eudicots. Publication of the *Amborella* genome (32) represented a significant advance for understanding the genetic basis of flowering plant origins. However, extant ‘basal’ angiosperms commonly exhibit habits ill-suited to maintaining large populations in controlled laboratory environments, long generation times [Figure 1A], and large genomes (> 1Gb) (62). *N. thermarum*, the smallest member of the Nymphaeaceae, has a short generation time of 4-5 months, has one of the smallest genomes described for any member of the Nymphaeales, and can both self-fertilize and outcross (31, 63, 64). A draft genome of *N. thermarum* represents a critical step in developing a system for functional genetics from within the basally-diverging angiosperm lineages — a tool that holds great promise for addressing Darwin’s “abominable mystery” regarding the early radiation of the most species-rich clade of plants on earth (65, 66).

## Methods

Exended methods are available in SI Appendix: Materials and Methods.

### Genome sample and sequence collection

Whole genome sequence data was collected from a live specimen of *Nymphaea thermarum* at the Arnold Arboretum at Harvard University. Genomic DNA was extracted using a modified CTAB extraction protocol (67). Short insert and mate-pair were prepared for sequencing using 1 ug DNA each with the Illumina TruSeq Library Kit and the Illumina Nextera Mate Pair Library Kit. Size selection was done using a Sage Science Pippin Prep machine with a target insert size of 3 kb. Both libraries were sequenced on an Illumina HiSeq 2500 sequencer with v4 chemistry (2×250 bp reads for the short insert library and 2×125 bp reads for the mate-pair library).

### Transcriptome sequence collection, assembly, and annotation

RNA was extracted from tissues (leaves, roots, floral buds, ovules) of multiple *N. thermarum* individuals grown at the Arnold Arboretum of Harvard University using a hot acid-phenol protocol (68). Libraries for RNA-Seq analysis were prepared by the Whitehead Institute Genome Technology Core, from total RNA (200-500ng) with the Apollo 324 system from WaferGen Biosystems using the WaferGen Prep-X Directional RNA-Seq kit according to manufacturer’s protocols to produce strand-specific cDNA libraries. Adapter-ligated cDNA fragments were enriched and amplified with 15 cycles of PCR using the HiFi NGS Library Amplification kit from KAPA Biosystems. Libraries with were multiplexed at equimolar concentration and sequenced on the Illumina HiSeq 2500 for 1×40 bases. rRNA reads were filtered out with Bowtie (69) by alignment to rRNA sequences from multiple *Nymphaeales.* Remaining reads were assembled using the Trinity pipeline (70, 71). TransDecoder was applied to contigs with the unstranded option to identify open reading frames (71). BLASTN, querying against the TAIR10 Arabidopsis cDNA database at an e-value of 0.05 was used to assign identities to the open reading frame encoded by the mRNA.

### Genome Assembly and Annotation

Reads from the short insert library were assembled into contigs using DISCOVAR *de novo* v52488 (72). We next used the mate-pair data and SSPACE v3.0 (73) to join contigs into scaffolds, which resulted in a draft *de novo* assembly of 368,014,730 bp in 6,225 scaffolds with an N50 of 275,242 bp (after removing scaffolds < 1 kb). Repeats were masked with RepeatMasker v4.0.5 (74). The quality of the draft genome was assessed with BUSCO v1.1(33) for conserved plant genes.

This assembly was then annotated using MAKER v2.31.8 (75) using protein evidence (protein sequences from select angiosperm reference genomes) and transcript evidence (transcripts of *N. thermarum).* Four iterations of MAKER were completed, each with annotations limited to > 20 amino acids on scaffolds > 5 kb in length. In the first run we initialized gene models for the *ab initio* software SNAP (76), and in subsequent runs the gene models were refined using the best ~2,500 genes and the ab initio program AUGUSTUS (77) was also added. MAKER generated 60,427 annotations, of which 25,760 had an annotation edit distance < 1, a Pfam domain (searched with InterProScan), or a blastp hit to the Uni/Swiss-Prot database. We used proteinortho v5.11 (78), with default settings, to identify shared and unique orthogroups between *Nymphaea thermarum, Arabidopsis thaliana*, and *Amborella trichopoda.*

### Phylogenomic analysis

We clustered homologs via an all-vs-all pairwise search with BLASTP v2.2.25 (79) with an e-value of 10^^^-20 followed by grouping with MCL v09-308 with an inflation value of 5.0 (80). We clustered in-paralogs at 98.5% identity using CD-HIT v4.6 (81) and retained the longest amino acid, or chose one randomly in case of ties. We required clusters to: 1) include at least four species with 2) at least one sequence from *Pinus* (for outgroup rooting), *Amborella*, and *Nymphaea* each, 3) include at least 100 amino acids for each sequence (82), 4) have a mean of less than five homologous sequences per species, and 5) have a median of less than two sequences per species (34). We aligned retained genes with Muscle v3.8.31 (83). We removed high-entropy regions of the alignment with TrimAl v1.2rev59 (84) and back-translated the amino acid alignment to codons using Pal2nal v14 (85). We calculated homolog trees on the back-translated codons using the GTRGAMMA model in RaxML v8.2.8 (86) with *Pinus* designated as the outgroup, running 100 rapid bootstraps, and selecting the best-scoring ML tree. We inferred orthologs following the “Maximal Occupancy” methods of Yang and Smith (87). Using only the inferred ortholog sequences, we then made new multiple alignments, filtered high-entropy regions, back-translated to codons, and calculated gene trees.

We concatenated the alignments into a supermaterix and identified parsimony informative sites using FASconCAT v1.02 (88). We calculated observed variability (OV) for every alignment position in the super matrix as described by Goremykin et. Al (89): for a given alignment position, the sum of all pairwise differences is divided by the total number of characters. We created two synthetic alignments for each gene cluster by partitioning parsimony-informative alignment positions into those evolving faster and slower than the median. We concatenated the fast and slow synthetic alignments into respective supermatricies. We calculated species trees on all three supermatricies and gene trees for the fast and slow evolving synthetic alignments using RaxML as described previously.We reconciled species trees from gene trees for all three rate categories using Astral-II v.4.10.6 (90) and MP-EST v1.4 (91).

### Phylogenomic dating, estimation of gene family expansion/contraction, and GO enrichment analysis

We used the ASTRAL II all-rate classes tree for phlyogenomic dating with r8s v1.5 (92). Four nodes were fixed or constrained, based on fossil evidence (5). Clusters used for phylogenomic analysis were filtered for a mean cluster size variance per taxa of less than or equal to 140, with a median size greater than or equal to 1, leaving a set of 8147 clusters included for further analysis with CAFÉ v3.1 (35).

For nodes of interest within the species tree, clusters for which cluster size changed significantly were identified (p-value of change from previous node < 0.05). Within a cluster of interest, the gene IDs of all *Arabidopsis thaliana* sequences were then used as the input for AgriGO Singular Enrichment Analysis (93) (against TAIR9 reference, Fischer test with Yekutieli multiple testing correction).

### Evolution of HD-ZIP III gene family

The cluster that contained HD-ZIP III members was defined by presence of *Arabidopsis thaliana* copies of *REVOLUTA, PHAVOLUTA, PHABULOSA, CORONA*, and *ATHB8.* This cluster was comprised of two clearly delineated sub-clades. One of these distinct sub-clades contained the HD-ZIP III genes; only this sub-clade was used in further gene family analysis. *Nuphar* putative homologs were identified using a HMMR v3.1b1(94) with aligned nucleic acid sequences of all *Nymphaea* and *Arabidopsis* homologs (aligned with MUSCLE v3.8.31 (83), manually trimmed to remove poorly conserved regions) against the *Nuphar* EST database (38) from the Ancestral Angiosperm Genome Project. Open reading frames were identified within the *Nuphar* sequences with Transdecoder v2.0.1 (71), and tBLASTx was used to identify *Nuphar* ORFs with sequence similarity to *Arabidopsis thaliana* HD-ZIP III genes. *Nuphar* sequences were added to the HD-ZIP III amino nucleic acid alignment, which was then used to estimate the gene family phylogeny with RAxML v8.2.8 (86) (GTRGAMMA model, 100 bootstrap replicates). Trees were viewed in FigTree v1.3.1 (95).

### Evolution of CLE peptides

Putative CLE peptide sequences were collected using a HMMR v3.1b1 (94) search of aligned nucleic acid sequences of all *Arabidopsis* homologs (aligned with MUSCLE v3.8.31 (83) and manually trimmed to remove the most poorly conserved regions) queried against a broad selection of plant genomes and transcriptomes. Amino acid sequences were aligned with MUSCLE v3.8.31 (83) and then trimmed to exclude the most poorly conserved regions. The resulting alignment of approximately 100 amino acids was used for CLANS (96) cluster analysis.

In order to normalize for different evolutionary distances represented by comparisons, the effect of evolutionary distance on cross-genera PAV was modeled in R v3.5.1, using the lm function and divergence times collected from TimeTree (97). The model was then used to remove the effect of evolutionary distance on PAV, to give APAV *(SI Appendix*, Figure S8). Taxa were classified by whether (VC) or not (NVC) they have a vascular cambium. For sequences for each CLE Groups of interest, APAV were sorted into VC-VC and NVC-VC categories depending on the taxa they originate from. Two-sample t-tests in R v3.5.1 were used to assess differences between APAV of the VC-VC and NVC-VC categories within each CLE subgroup. APAV were also compared between the sets of all sequences from taxa or clades of interest and the set of all sequences from VC taxa.

### Microscopy

*N. thermarum* rhizomes older than 1 year were collected from specimens at the Arnold Arboretum of Harvard University, embedded in JB4 resin (Electron Microscopy Sciences), and processed for microscopy (98). Sections were stained with periodic acid-Schiff (PAS) reagent and toluidine blue (99). Prepared slides of *Medicago truncatula* and *Zea mays* stems were purchased from Triarch incorporated (Ripon, WI, USA). Bright-field and differential interference contrast images were recorded with a Zeiss Axio Imager Z2 microscope equipped with a Zeiss HR Axiocam digital camera (Zeiss, Oberkochen, Germany).

## Data Availability

Raw sequence data, whole-genome assembly, and transcriptomes of *N. thermarum* have been submitted to the National Center for Biotechnology Information (NCBI) database under BioProject <u>PRJNA508901</u>. Biological material and all other data are available as Extended or Supplemental Data, or from the corresponding authors upon request.

## Acknowledgements

We acknowledge support from the National Science Foundation: IOS-0919986 awarded to W.E.F., DEB-1120243 awarded to C.C.D., MCB-1453459 awarded to M.G., IOS-1416825 awarded to S.M., and IOS-1812116 awarded to R.A.P.. Botanische Gärten der Universität Bonn provided original plant material for propagation.

## Author Contributions

M.G.,W.E.F., and C.C.D. conceived of original premise of the project. R.A.P. grew plant samples and extracted genomic DNA. M.M. and J.J. collected and processed plant material for RNA-seq. P.R.V.S. performed assembly and annotation of transcriptome data. J.M.D-C. processed samples for genomic DNA library preparation and performed genome assembly and annotation, using methods developed with S.M.. Z.X. and C.G. performed ortholog clustering and phylogenomic analysis. C.G. and R.A.P. performed gene-family contraction/expansion analysis. R.A.P. performed analysis of HD-ZIP III and CLE gene families. R.A.P. collected, processed, and sectioned rhizome material, and imaged prepared slides. R.A.P. wrote the manuscript with input from M.G., S.M., C.C.D., and W.E.F.

## Author Information

Reprints and permissions information is available at www.nature.com/reprints. The authors declare no competing financial interests. Correspondence and requests for materials should be addressed to W.E.F. and C.C.D.

