## Supplementary Information Appendix for "Water lily (*Nymphaea thermarum*) draft genome reveals variable genomic signatures of ancient vascular cambium losses"

***SI Figures***

***Figure S1___________________________page 2***

***Figure S2___________________________page 3***

***Figure S3___________________________page 4***

***Figure S4___________________________page 5***

***Figure S5___________________________page 6-7***

***Figure S6___________________________page 8***

***Figure S7___________________________page 9***

***Figure S8___________________________page 10***

***SI Tables***

***Tables S1-S3 (captions)_______________page 11***

***Table S4 (caption and table)__________page 11***

***Supplementary Materials and Methods_______page 12-24***

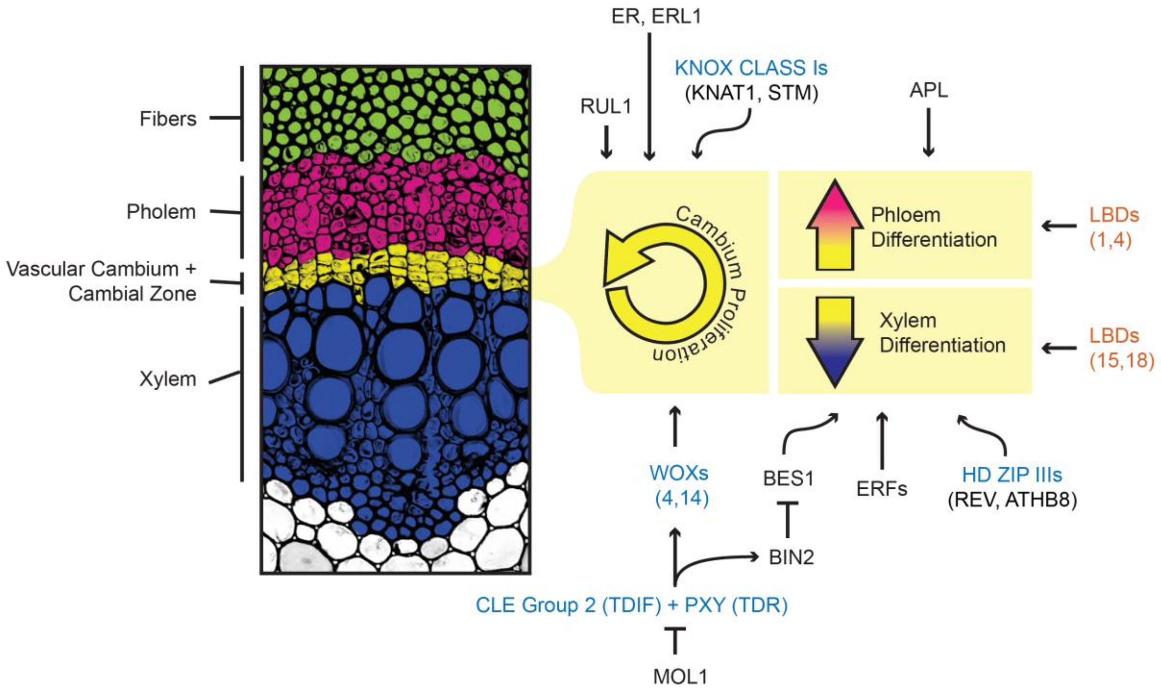


***SI***

***Appendix,* Figure S1:** Summary of regulation of secondary vascular tissue development in flowering plants. As a meristematic tissue, the vascular cambium (yellow) becomes active and undergoes cell divisions. The resulting cells can participate in cambium proliferation, abaxial differentiation of phloem tissue (pink), or adaxial differentiation of xylem tissue (blue). Genes known to regulate these processes during secondary vascular development are shown. Color of text indicates which species gene function experiments have been performed in: black = *Arabidopsis thaliana*, orange = *Populus trichocarpa*, blue = multiple flowering plant species (*A. thaliana*, *P. trichocarpa*, and/or *Zinnia elegans*). When multiple species have been studied for a gene, names are given for *A. thaliana* orthologs.

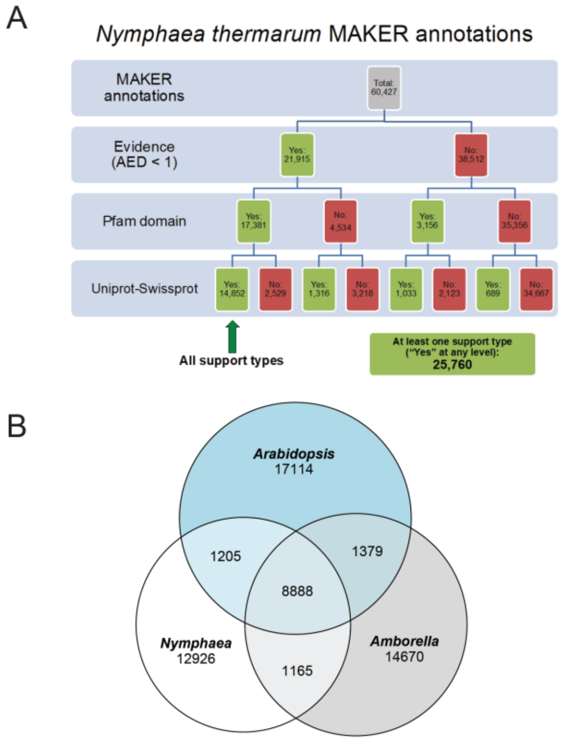


***SI Appendix,* Figure S2:** Annotation of the draft genome of *Nymphaea thermarum*. A) Quality control for MAKER annotations of *N. thermarum* draft genome assembly. Three types of evidence were used to confirm MAKER annotations: AED (Annotation Edit Distance) score < 1, presence of a Pfam domain, and/or a blastp hit against the full UniProt-SwissProt protein database. 25,760 annotations had at least one type of support; 14,852 were supported by all three types. B) Ortholog analysis of annotated proteins from the genomes of *Amborella trichopoda* (n=26,846), *Arabidopsis thaliana* (35,386), and *N. thermarum* (n=25,760). Each ortholog group could include 1-3 species and 1-n proteins; species could have multiple proteins that are assigned to the same cluster due to high similarity. *N. thermarum* is represented in 24,184 ortholog groups, and in 150 (0.6%) of these there are >1 proteins. The low number of ortholog groups with multiple *N. thermarum* copies indicates that there is little pseudo-replication of regions due to poor quality assembly.

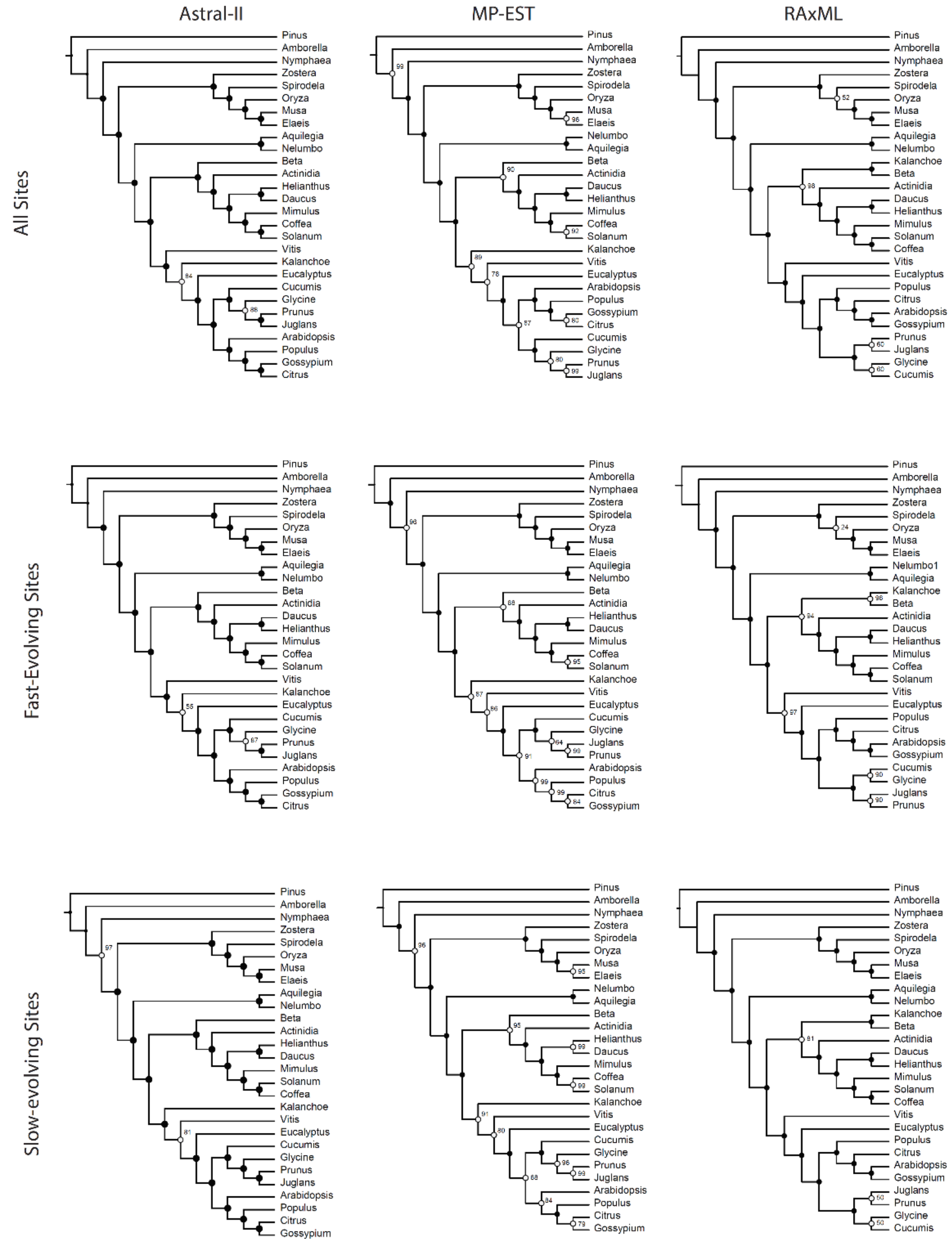


***SI Appendix,*** **Figure S3:** Results of phylogenomic analyses. Reconciled species trees (ASTRAL-II, MP-EST ) and calculated species trees (RAxML) for three supermatrices (all sites, fast-evolving site, or slow-evolving sites), which represent data from 1,439 orthologs clusters from 28 species. Support values represent multilocus bootstrap support (ASTAL-II), jackknife support (MP-EST), or bootstrap support (RAxML). Black dots indicate support values = 100, white dots indicate support values <100 (values reported at nodes).

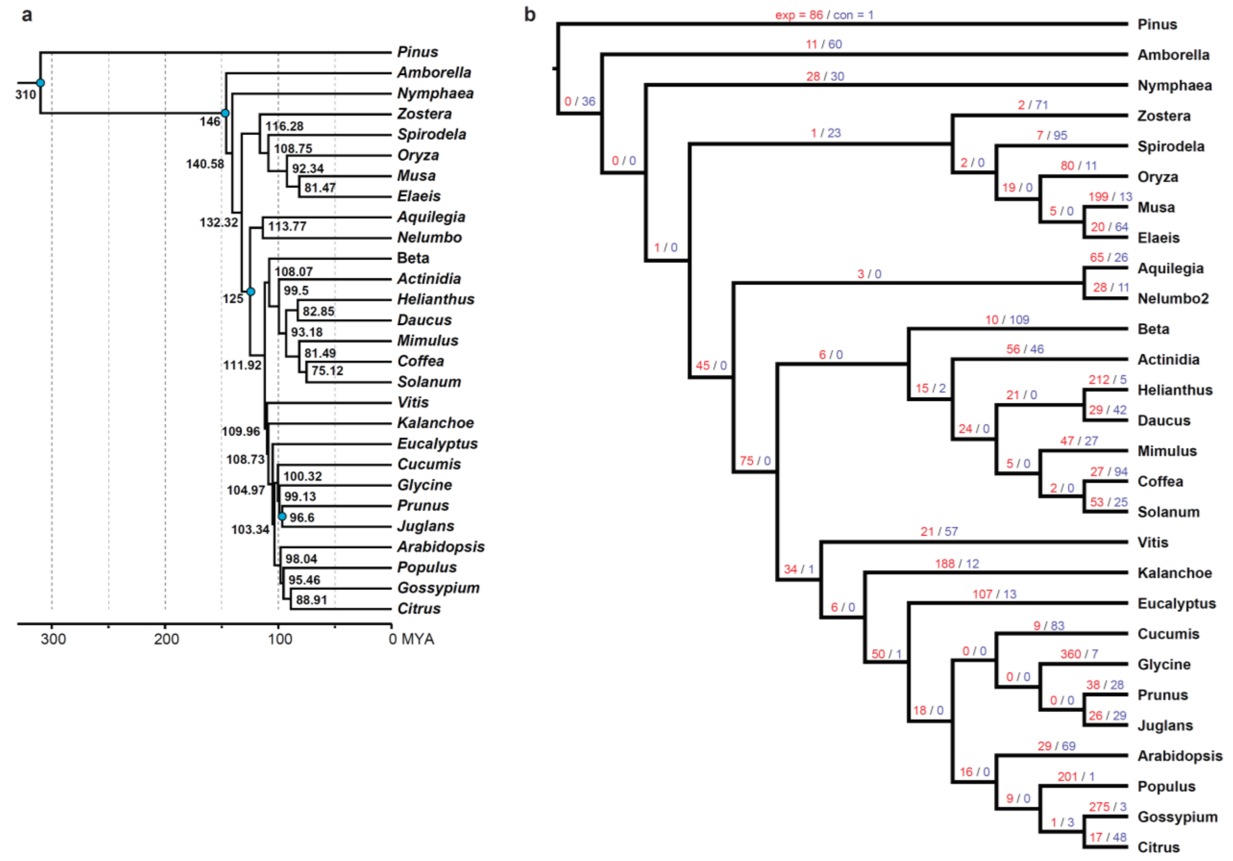


***SI Appendix,*** **Figure S4:** *SI Appendix, Figure S* 4: Summary of divergence age estimates and CAFÉ analysis for significant gene family expansion and contraction. A) Fossil-calibrated phylogenomic analysis. Blue dots indicate age and placement of fossils or other estimates used for calibration.  B) For each branch, the number of significantly expanded (red, “exp”) and contracted (purple, “con”) gene clusters is reported.

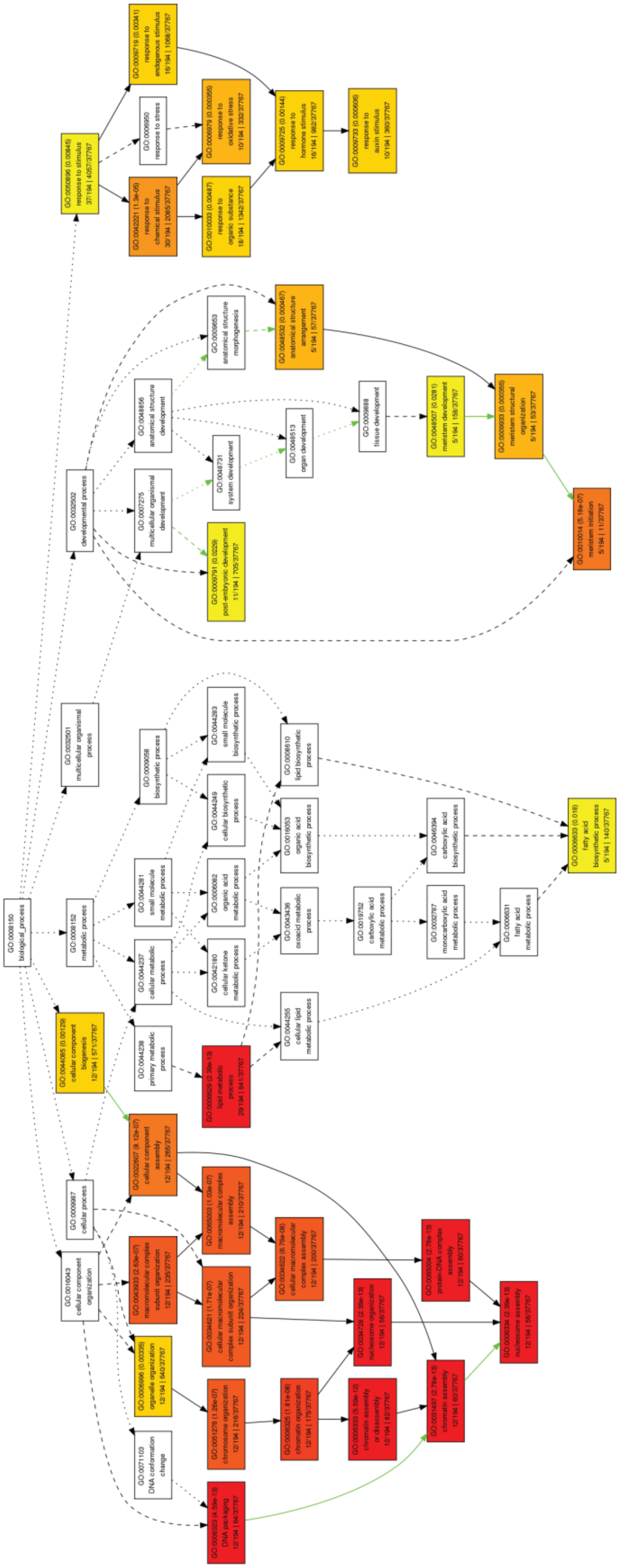

***(Preceding Page) SI Appendix,*** **Figure S5:** Singular GO term enrichment analysis for the set of all gene clusters significantly contracted in *Nymphaea thermarum*. Relationships between GO terms are displayed with arrows (black = hierarchical relationship, green = negative relationship, large dash = two significant nodes, small dash = one significant node), and box colors indicate level of significance of enrichment (red to yellow = very significant to significant, white = no significant enrichment). Each box reports GO ID (top left), p-value for enrichment (top right), number of genes within the test dataset that are annotated with the GO term (bottom left), and number of genes within reference dataset that are annotated with the GO term (bottom right). Root terms are at the top of the diagram, child terms towards the bottom. Diagram created with AgriGO v1.2.

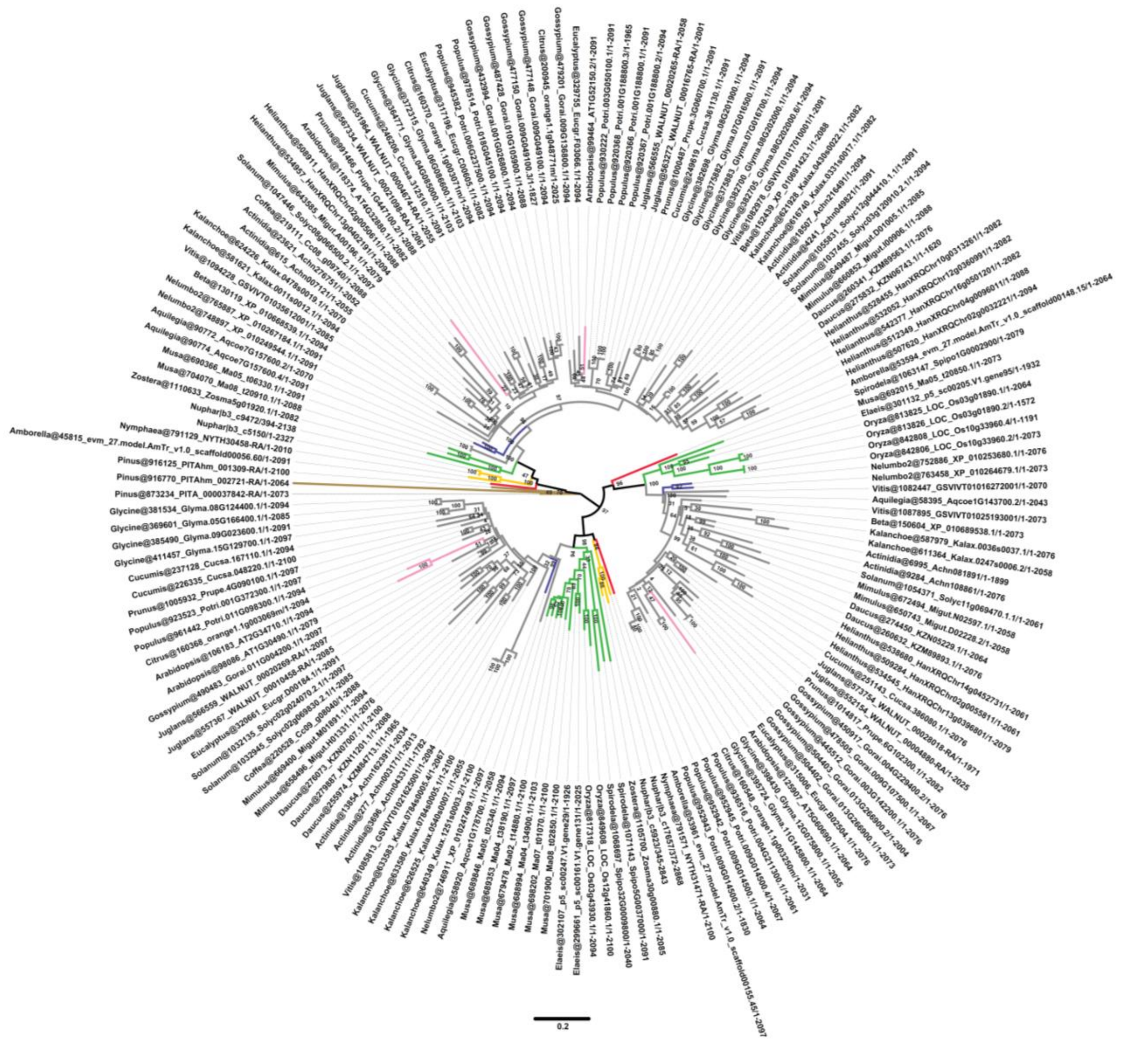


***SI Appendix,*** **Figure S6:** Detailed HD-ZIP III gene family phylogeny. Branches are color-coded similarly to Figure 2C. Sequence names include both name using during analysis (before "_") and gene ID from respective genome annotations (after "_"). Branch length (scale at bottom) represents changes per site, numbers at nodes indicate bootstrap support values.

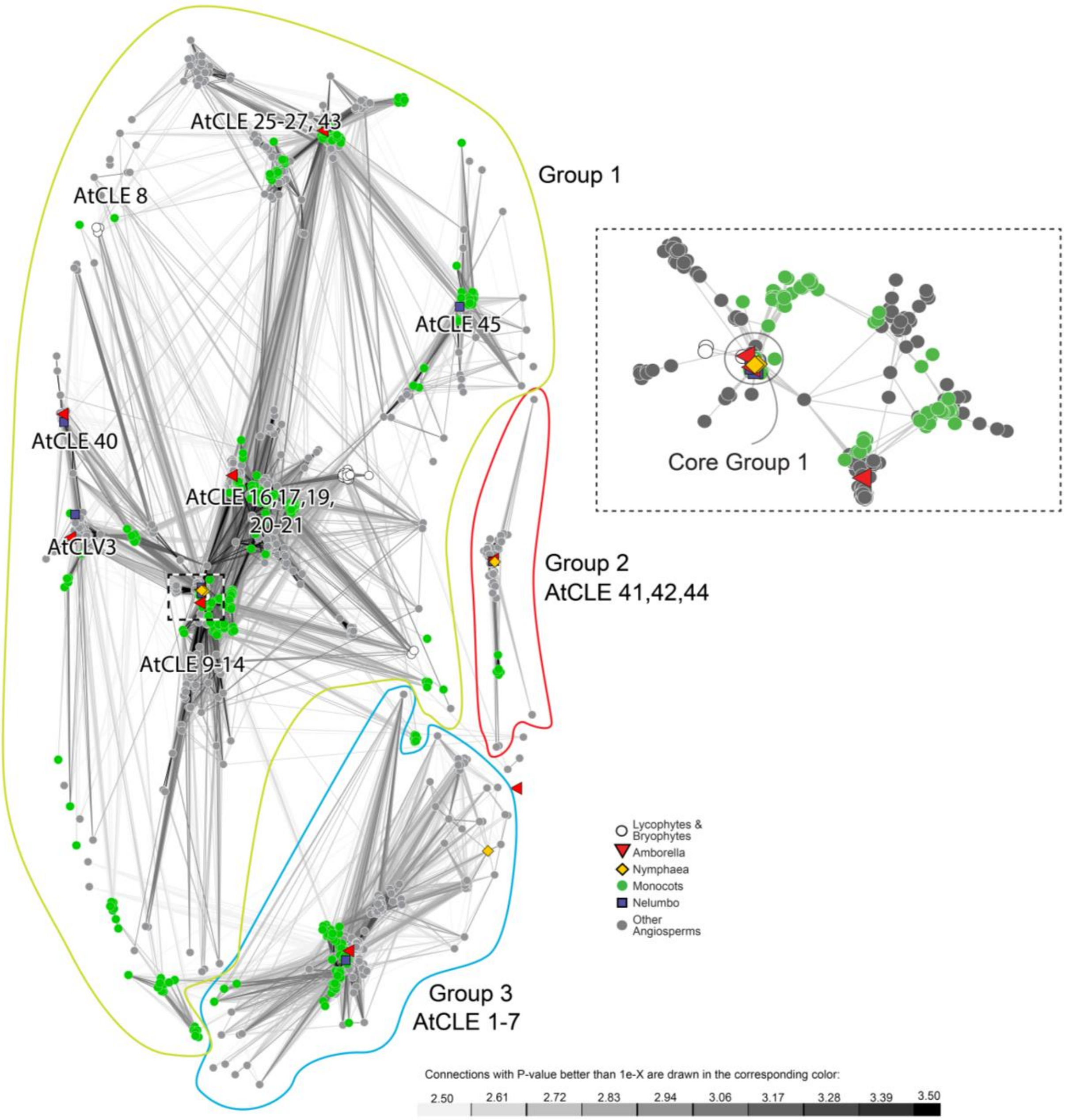


***SI Appendix,*** **Figure S7:** 2-dimensional CLANS clustering of conserved amino acid sequences of CLE peptides in land plants. Dots/shapes represent individual sequences, coded to indicate whether they are from species within clades that have vascular cambia (VC) or do not have vascular cambia (NVC). Clustering was performed using connections with P-values less than 5e-6; connections with P-values less than 1e-3 are shown. Groups or subgroups with Arabidopsis CLV3 and CLE sequences are indicated. Box with black/white outline indicates area considered for selection of Core Group 1, separate clustering of these sequences are shown in the inset (connections with P-values less than 1e-7 were used and are shown). Circled area indicates sequences used in Core Group 1.

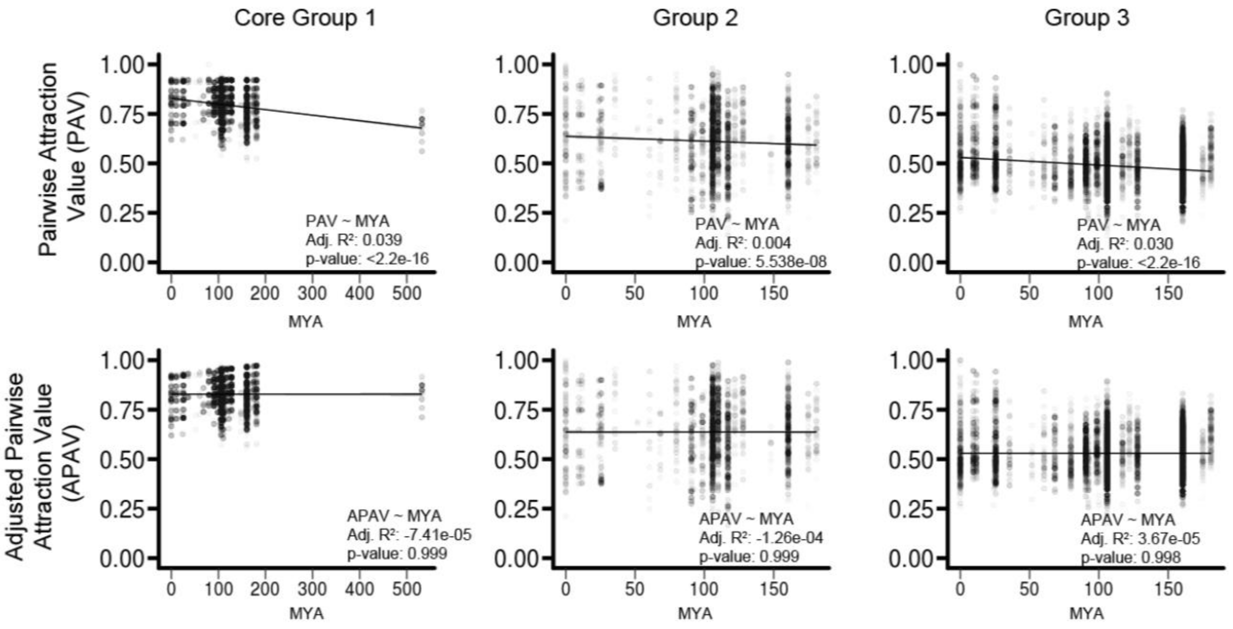


***SI Appendix,*** **Figure S8:** Modeling effect of evolutionary distance (time since divergence in millions of years, MYA) on Pairwise Attraction Values (PAV). Each data point represents the similarity (PAV) of two CLE sequences from different taxa that resulted from CLANS clustering analysis; the highly conserved resides at the C terminus were included in the analysis. Adjusted R-squared and p-value are reported for a linear model of the effect of time since divergence on PAV or Adjusted PAV (APAV). For each of three sets of CLE genes (Core Group 1, Group 2, and Group 3), a small but statistically significant negative relationship between time since the two taxa diverged and CLE sequence similarity is shown (top row). The effect of time since divergence was removed to give APAV. There is no significant relationship between APAV and time for any of the three groups of CLE genes (bottom row).

***SI Appendix,*Table S1:** Information on genome assembly and annotation [downloadable file]

***SI Appendix,*Table S2:** List of significantly contracted and expanded clusters in *Nymphaea thermarum*, *Nelumbo nucifera*, and the 'stem' branch to extant monocots, with GeneIDs and annotation information for *Arabidopsis thaliana* homologs present in each cluster. [downloadable file]

***SI Appendix,*Table S3:** Full result of Singular GO Enrichment Analysis for sets of gene clusters expanded (Exp.) or contracted (Con) in sets of cambium-less angiosperm lineages (*Nymphaea*, stem branch to extant monocots, *Nelumbo*). [downloadable file]

***SI Appendix,*Table S4:** Comparisons of CLE peptide sequence similarity between species with (VC) and without (NVC) a vascular cambium

| | | | | **2-sample t-test (VC-VC vs. NVC-VC)** | | |
|---|---|---|---|---|---|---|
| **CLE Group** | Category | n | mean APAV | *t*-value | df | *p*-value |
| **1, core** | VC-VC | 9998 | 0.829716 | 4.4201 | 6257.9 | 0.00001003 |
| | NVC-VC | 3500 | 0.8232659 | | | |
| **2** | VC-VC | 7396 | 0.6415516 | 12.789 | 645.19 | < 1E-16 |
| | NVC-VC | 522 | 0.5693744 | | | |
| **3** | VC-VC | 16256 | 0.5265861 | -6.3938 | 25006 | 1.647E-10 |
| | NVC-VC | 11008 | 0.534508 | | | |

**Supplementary Materials and Methods:**

**Genome sample and sequence collection**

We collected whole genome sequence data from a live specimen of *Nymphaea thermarum* (ID= Rp0033) at the Arnold Arboretum at Harvard University. Genomic DNA was extracted using a modified CTAB extraction protocol(1) from leaf, root, and bud tissues. Briefly, 0.5 g tissue was ground with liquid nitrogen, to which 4ml of CTAB buffer (to which 0.2 g Polyvinylpyrrolidinone (PVP 40) and 100 µl 2-Mercaptoethanol had been added) and then incubated for 1 hour at 65 ̊ C. Samples were treated with RNase A for 15 minutes at 37 ̊ C, and then a chloroform extraction was performed twice. DNA was precipitated with cold isopropanol, resuspended in TE buffer with 0.5 volumes of 7.5 M ammonium acetate, precipitated with cold 100% ethanol, washed with cold 70% ethanol, and then resuspended in nuclease-free water. We assessed extract quality with fluorometric quantitation (Invitrogen Qubit) and electrophoresis (Agilent Tapestation). All extracts had sufficient concentration and proportion of high molecular weight fragments for library preparations, so we pooled extracts before the collection of whole genome sequence data. We then prepped two different libraries for Illumina sequencing: short insert and mate-pair. For the short insert library we used the Illumina TruSeq Library Preparation Kit with 1 µg of input genomic DNA, following the manufacturer's protocol. The final purified library was evaluated with an Agilent Tapestation assay, which showed a distribution of fragment sizes with a mode of ~500 bp and no signs of adapter dimer contamination. For the mate-pair library we used the Illumina Nextera Mate Pair Library Preparation Kit with 1 µg of input genomic DNA and followed the manufacturer's protocol. Size selection was done using a Sage Science Pippin Prep machine with a target insert size of 3 kb, however, an Agilent Tapestation assay of the post-size selection product showed a peak at ~ 5 kb. Both libraries were sequenced on an Illumina HiSeq 2500 sequencer with v4 chemistry. We collected 2x250 bp reads for the short insert library and 2x125 bp reads for the mate-pair library.

**Transcriptome sequence collection, assembly, and annotation**

RNA was extracted from tissues (leaves, roots, floral buds, ovules) of multiple *N. thermarum* individuals (from the Arnold Arboretum of Harvard University) using a hot acid-phenol protocol(2). Libraries for RNA-Seq analysis were prepared by the Whitehead Genome Technology Core, from total RNA (200-500ng) with the Apollo 324 system from WaferGen Biosystems using the WaferGen Prep-X Directional RNA-Seq kit according to manufacturer's protocols to produce strand-specific cDNA libraries. Adapter-ligated cDNA fragments were enriched and amplified with 15 cycles of PCR using the HiFi NGS Library Amplification kit from KAPA Biosystems. Libraries were quantified by qPCR using the KAPA Biosystems Illumina Library Quantification kit according to kit protocols. Libraries with distinct TruSeq indexes were multiplexed by mixing at equimolar concentration and sequenced on the Illumina HiSeq 2500 for 40 bases in single-read, high-output mode. Image processing and base calling were preformed using the standard Illumina pipeline. Reads were trimmed and filtered to improve quality using "*fastq_quality_trimmer -t 34 -l 47"*. Only paired end reads were retained and single reads were discarded. rRNA reads were filtered out with Bowtie(3) by alignment, allowing three mismatches, to rRNA sequences from multiple *Nymphaeles* species including *Nymphaea leibergii*, *Nymphaea tetragona*, *Nymphaea alba*, *Nymphaea lingulata*, *Nymphaea rudgeana*, *Nuphar variegata, Nuphar pumila, Nuphar shimadai, Nuphar advena, Cabomba caroliniana* and *Nymphaea immutabilis.* After removal of rRNA sequences, 84.5 million reads that passed these quality controls were assembled into a 58,177 contig transcriptome using the Trinity pipeline(4, 5) without normalization. TransDecoder was applied to these contigs with the unstranded option to identify open reading frames(5). 27,115 transcripts with an N50 of 1383 bp were identified. BLASTN, querying against the TAIR10 Arabidopsis cDNA database at an e-value of 0.05 was used to assign identities to the open reading frame encoded by the mRNA.

**Genome Assembly and Annotation**

We collected 44,640,690 paired reads for the short insert library (~40x coverage of estimated genome size of 498,780,000 bp based on flow cytometry) that passed the initial Illumina quality filter, and assembled these reads into contigs using DISCOVAR *de novo* v52488(6). Ignoring contigs < 1 kb in length, the DISCOVAR *de novo* assembly contained 363,660,557 bp in 22,407 contigs with an N50 of 50,076 bp. We next used the mate-pair data (49,345,536 paired reads; ~20x coverage) and SSPACE v3.0(7) to join contigs into scaffolds, which resulted in a draft *de novo* assembly of 368,014,730 bp in 6,225 scaffolds with an N50 of 275,242 bp (after removing scaffolds < 1 kb). Repeats in the assembly were then masked with RepeatMasker v4.0.5(8). The quality of the draft genome was assessed with BUSCO v1.1(9), in which we searched for a set of 956 genes that are conserved across plants. The analysis recovered 865 (90%) plant BUSCOs as complete copies, and another 35 (4%) as fragmented.

This assembly was then annotated using MAKER v2.31.8(10). For protein evidence we used a set of 160,490 protein sequences from reference genomes of *Amborella trichopoda*, *Solanum lycopersicum*, *Arabidopsis thaliana*, and *Zea mays*, and from the UniProt database for "basal Magnoliophyta." For transcript evidence we used the transcriptome assembly (27,115 transcripts) of *N. thermarum* derived from a variety of tissue types. Four iterations of MAKER were completed, each with annotations limited to > 20 amino acids on scaffolds > 5 kb in length. In the first run we initialized gene models for the *ab initio* software SNAP(11), and in subsequent runs the gene models were refined using the best ~2,500 genes and the ab initio program AUGUSTUS(12) was also added. MAKER generated 60,427 annotations, of which 25,760 had an annotation edit distance < 1, a Pfam domain (searched with InterProScan), or a blastp hit to the Uni/Swiss-Prot database. We used proteinortho v5.11(13), with default settings, to identify shared and unique orthogroups between *Nymphaea thermarum*, *Arabidopsis thaliana*, and *Amborella trichopoda*.

**Phylogenomic analysis**

We clustered homologs via an all-vs-all pairwise search with BLASTP v2.2.25(14) with an e-value of 10^-20 followed by grouping with MCL v09-308 with an inflation value of 5.0(15). We clustered in-paralogs at 98.5% identity using CD-HIT v4.6(16) and retained the longest amino acid, or chose one randomly in case of ties. We required clusters to: 1) include at least four species with 2) at least one sequence from *Pinus* (for outgroup rooting), *Amborella*, and *Nymphaea* each, 3) include at least 100 amino acids for each sequence(17), 4) have a mean of less than five homologous sequences per species, and 5) have a median of less than two sequences per species(18). We aligned retained genes with Muscle v3.8.31(19). We removed high-entropy regions of the alignment with TrimAl v1.2rev59(20) "-automated1 -colnumbering" and back-translated the amino acid alignment to codons using Pal2nal v14(21). We calculated homolog trees on the back-translated codons using the GTRGAMMA model in RaxML v8.2.8(22) with *Pinus* designated as the outgroup, running 100 rapid bootstraps, and selecting the best-scoring ML tree. We inferred orthologs following the "Maximal Occupancy" methods of Yang and Smith(23): we masked monophyletic tips, retained homolog trees with monophyletic *Pinus* sequences, and trimmed paralogs from root to tip. Using only the inferred ortholog sequences, we then made new multiple alignments, filtered high-entropy regions, back-translated to codons, and calculated gene trees.

We concatenated the alignments into a supermaterix and identified parsimony informative sites using FASconCAT v1.02(24). We calculated observed variability (OV) for every alignment position in the super matrix as described by Goremykin et. al(25): for a given alignment position, the sum of all pairwise differences is divided by the total number of characters. We created two synthetic alignments for each gene cluster by partitioning parsimony-informative alignment positions into those evolving faster and slower than the median. We concatenated the fast and slow synthetic alignments into respective supermatricies. We calculated species trees on all three supermatricies and gene trees for the fast and slow evolving synthetic alignments using RaxML as described previously.

We reconciled species trees from gene trees for all three rate categories using Astral-II v.4.10.6(26) and MP-EST v1.4(27). We ran Astral-II with multi-locus bootstrapping using the RaxML rapid bootstraps as input. We estimated a jackknife branch support(28) for the MP-EST species tree by running the program 100 times, each with a unique 10% subset of randomly chosen gene trees. We used the GTRGAMMA model in RaxML to project branch lengths onto the coalescent tree topologies using the concatenated supermatricies.

**Phylogenomic dating**

We used the ASTRAL II all-rate classes tree for phlyogenomic dating with r8s v1.5(29). Four nodes were fixed or constrained, based on fossil evidence(30): stem angiosperms fixed at 310 MYA, crown eudicot minimum of 125, and Rosales + Fagaceae minimum of 96.6. Additionally, crown angiosperms were constrained with a minimum of 136 and maximum of 146 MYA(30, 31). For analysis with r8s, the divtime method used was the PL method with TN algorithm, penalty was set to additive, and smoothing was set to 0.0032 (based on an initial run with cross validation, crossv=yes cvstart=-2 cvinc=0.1 cvnum=30).

**Estimation of gene family expansion/contraction and GO enrichment analysis**

Clusters used for phylogenomic analysis were filtered for a mean cluster size variance per taxa of less than or equal to 140, with a median size greater than or equal to 1, leaving a set of 8147 clusters included for further analysis with CAFÉ v3.1(32). For the CAFÉ analysis, a two-parameter model was applied for the time-calibrated r8s output tree; terminal branches for *Glycine* and *Pinus* were assigned to one parameter class and all other taxa and internal branches to another (reflecting the substantially larger sizes of the *Glycine* and *Pinus* genomes). The analysis resampled 1000 times to calculate p-values.

For nodes of interest within the species tree, clusters for which cluster size changed significantly were identified (p-value of change from previous node < 0.05). Within a cluster of interest, the gene IDs of all *Arabidopsis thaliana* sequences were then used as the input for AgriGO Singular Enrichment Analysis(33) (against TAIR9 reference, Fischer test with Yekutieli multiple testing correction).

**Evolution of HD-ZIP III gene family**

The cluster that contained HD-ZIP III members was defined by presence of *Arabidopsis thaliana* copies of *REVOLUTA*, *PHAVOLUTA*, *PHABULOSA*, *CORONA*, and *ATHB8*. This cluster, however, was comprised of two clearly delineated sub-clades. One of these distinct sub-clades contained the HD-ZIP III genes; only this sub-clade was used in further gene family analysis. *Nuphar* putative homologs were identified using a HMMR v3.1b1(34) search (evalue-cutoff = 1e-10) with aligned nucleic acid sequences of all *Nymphaea* and *Arabidopsis* homologs (aligned with MUSCLE v3.8.31(19), manually trimmed to remove poorly conserved regions) against the *Nuphar* EST database(35) from the Ancestral Angiosperm Genome Project. Open reading frames (ORFs) were identified within the *Nuphar* sequences with Transdecoder v2.0.1(5), and tBLASTx was used to identify *Nuphar* ORFs with sequence similarity to *Arabidopsis thaliana* HD-ZIP III genes using a e-value cut-off of 1e10. Four *Nuphar* sequences were identified, translated, and added to the HD-ZIP III amino nucleic acid alignment, which was then used to estimate the gene family phylogeny with RAxML v8.2.8(22) (GTRGAMMA model, 100 bootstrap replicates). Trees were viewed in FigTree v1.3.1(36).

**Evolution of CLE peptides**

Due to relatively low conservation among CLE sequences, as well as their short sequence length, a CLANS(37) clustering approach was taken for characterizing patterns of divergence within this gene family. Putative CLE peptide sequences were collected using a HMMR v3.1b1(34) search of aligned nucleic acid sequences of all *Arabidopsis* homologs (aligned with MUSCLE v3.8.31(19) and manually trimmed to remove the most poorly conserved regions) queried against the following databases: annotated peptide sequences of 47 Phytozome v12(38) genomes, genomes of *Nymphaea thermarum* and *Nelumbo nucifera*(39), available angiosperm transcriptomes from the OneKP initiative(40), and *Nymphaea thermarum* transcriptomes. Amino acid sequences were aligned with MUSCLE v3.8.31(19) and then trimmed to exclude the most poorly conserved regions. The resulting alignment of approximately 100 amino acids was used for CLANS cluster analysis (BLOSUM62 scoring matrix, HSP cut-off 1e-4, run for >10,000 iterations, singletons removed). The resulting clusters (subgroups) were similar to those that have been previously reported, based on membership of *Arabidopsis thaliana* and *Oryza sativa* homologs(41, 42).

Pairwise attraction values (PAV, rescaled so that lowest value = 0 and highest value = 1) between all taxa were extracted for subgroups of interest: Group 2, Group 3, and a selection of 'core' Group 1. A subset of group Group 1 was used because Group 1 is much more broadly defined than the other groups, and in fact is comprised of several smaller, distinct subgroups(42). In order to normalize for the different evolutionary distances represented by comparisons, the effect of evolutionary distance on PAV was modeled. For all cross-genera comparisons, the estimated divergence time between genera was collected from TimeTree(43). In R v3.5.1, the lm function was used to model the effect of evolutionary distance (divergence time) on PAV (model = PAV~evolutionary distance); fit of the model with evolutionary distance as a fixed effect was a significant improvement over fit of the null model. The slope of the linear model was then used to remove the effect of evolutionary distance on PAV, to give the Adjusted Pairwise Attraction Value (Adjusted PAV, APAV) (Supp Figure 8). Taxa were classified by whether or not they have a vascular cambium (VC = have a vascular cambium, NVC = do not have a vascular cambium), and then within each subgroup of interest, pairwise comparisons and their Adjusted PAV were sorted into VC-VC and NVC-VC categories. Two-sample *t*-tests in R v3.5.1 were used to assess differences between APAV of the VC-VC and NVC-VC categories within each CLE subgroup. APAV were also compared between the sets of all sequences from taxa or clades of interest (*Amborella trichopoda*, *Nymphaea thermarum*, Magnoliids, monocots, *Nelumbo nucifera*, Core Eudicots) and the set of all sequences from VC taxa.

**Microscopy**

*N. thermarum* rhizomes older than 1 year were collected from specimens at the Arnold Arboretum of Harvard University, and fixed in 4 % v/v acrolein (Polysciences, New Orleans, LA, USA) in 1× PIPES buffer pH 6.8 (50 mm PIPES, 1 mm MgSO4, 5 mm EGTA) for 24 h. Fixed material was then rinsed three times (1 hour each time) with 1× PIPES buffer, dehydrated through a graded ethanol series and stored in 70 % ethanol. Samples were then dehydrated though a graded ethanol series up to 100 % ethanol, then infiltrated with and embedded in glycol methacrylate (JB-4 Embedding Kit, Electron Microscopy Sciences, Hatfield, PA, USA). Embedded materials were serially sectioned in 2-µm thick ribbons with a Leica RM2155 rotary microtome and mounted onto slides. Sections were stained with periodic acid–Schiff (PAS) reagent for insoluble polysaccharides and counterstained with toluidine blue for general tissue structure(44). Prepared slides of *Medicago truncatula* and *Zea mays* stems were purchased from Triarch incorporated (Ripon, WI, USA). Bright-field and differential interference contrast images were recorded with a Zeiss Axio Imager Z2 microscope equipped with a Zeiss HR Axiocam digital camera (Zeiss, Oberkochen, Germany).

**Data Availability**

Raw sequence data, whole-genome assembly, and transcriptomes of *N. thermarum* have been submitted to the National Center for Biotechnology Information (NCBI) database under BioProject PRJNA508901. Biological material and all other data are available as Extended or Supplemental Data, or from the corresponding authors upon request.
